# Regulator of G-Protein Signaling 2 Knockout in CD4+ T Cells Promotes Anti-Inflammatory T Cells, Enhancing Ovulation, and Oocyte Yield

**DOI:** 10.1101/2024.10.15.618561

**Authors:** Marika Raff, Trinity Benton, Daniel Brummond, Denali Kovach, Olivia Bunton, Ella Janky, Eyup Hakan Duran, Douglas G. Scroggins, Gabrielle Gray, Sabrina Marie Scroggins

## Abstract

**Objective:** To determine the downstream effects on ovarian function and immune cell differentiation in the ovary and uterus using a model in which RGS2 was knocked out specifically in CD4+ T cells.

**Design:** Laboratory based experiments with female mice.

**Animals:** Female congenic (fully backcrossed) and non-congenic (mixed strain) mice with CD4 T cell-specific RGS2 knockout.

**Exposure:** Four-week-old female CD4 RGS2 knockout (CD4 RGS2^KO^) mice and their littermate controls (CD4 RGS2^CTL^) were subjected to superovulation using pregnant mare serum gonadotropins.

**Main Outcome Measures:** Oocyte numbers, lymphocyte populations in the ovary and uterus, and serum estradiol and progesterone concentrations.

**Results:** In non-congenic (mixed strain) mice, CD4 RGS2 knockout (KO) promoted higher oocyte ovulation and increased uterine total leukocyte numbers. Similarly, congenic (fully backcrossed strain) mice showed higher oocyte numbers and increased uterine total leukocytes in the CD4 RGS2^KO^ mice compared to CD4 RGS2^CTL^ mice. Pro-inflammatory CD4+ T helper (T_H_) 1 and T_H_17 cell frequencies in the ovary and uterus were unchanged, while Treg and T_H_2 cell frequencies were elevated, along with increased concentrations of estradiol and progesterone in the serum of CD4 RGS2^KO^ mice.

**Conclusion:** Our study highlights the important role of RGS2 in CD4+ T cells within the context of reproduction. The dysregulation of immune responses due to RGS2 knockout in CD4+ T cells appears to enhance oocyte production. Further research is warranted to elucidate the precise mechanisms by which RGS2 influences reproductive outcomes, including its impact on fecundability, endometrial receptivity, and successful implantation.

## Introduction

Infertility affects 10-15% of couples, posing emotional and financial burdens. For some, the cause is identifiable, but for others, it remains unexplained. Successful reproduction requires ovarian follicular development, ovulation, mature oocytes, embryo development, and a receptive endometrium. Understanding the physiological processes involved in reproduction is key to identifying and treating lesser-known causes of infertility. The immune system’s role in reproduction is gaining attention as its involvement in these processes becomes clearer.

The immune system regulates inflammation, angiogenesis, and cytokine release, all essential for follicular growth, ovulation, endometrial maintenance, and embryo implantation. However, dysregulation of immune responses can impair reproductive function. Immune cells within the ovary and uterus orchestrate inflammation and tissue remodeling necessary for ovulation and implantation [1-4]. G-protein-coupled receptors (GPCRs) control these immune processes, and their signaling is regulated by proteins like RGS2 [5], which has critical roles in both oocyte and immune cell function [1-4, 6-9] (Supplementary Figure 1).

RGS2 is expressed in various tissues [1-9], including the ovary and uterus, where it plays roles in embryo implantation, vascular adaptations, and ovulation [4, 8-10]. In the ovary, RGS2 prevents premature oocyte activation and is involved in meiotic spindle formation and chromosome segregation [2, 7]. Additionally, RGS2 has been found as a potential biomarker associated with oocyte developmental competence and clinical pregnancy [11]. It also regulates immune responses, influencing T cell proliferation and cytokine production [6]. Studies suggest that RGS2’s regulation of GPCR signaling is crucial for reproductive tissues and immune function.

Immune cells are vital for reproductive events such as follicular development, ovulation, and corpus luteum function as well as endometrial development and maintenance. In the ovary, immune cells such as T cells, B cells, and macrophages contribute to tissue remodeling and inflammation. During ovulation, these cells increase in number [1], facilitating follicular rupture and oocyte release [12]. In the endometrium, immune cells are predominantly tissue-resident, with T cells playing a role in endometrial maintenance and immune tolerance during implantation and pregnancy. Dysregulated immune responses, particularly involving T regulatory cells (Tregs), are linked to implantation failure and pregnancy loss [13].

While individual roles of RGS2 and CD4+ T cells in reproduction are known, the impact of RGS2 loss specifically in CD4+ T cells on ovarian and uterine function remains unexplored. We hypothesized that loss of RGS2 in CD4+ T cells would reduce inflammatory T helper (T_H_) cells in reproductive tissues, impair ovulation, and decrease oocyte yield.

## Material and Methods

### Animal Studies

All animal procedures were approved by the Institutional Animal Care and Use Committees (IACUC) of the University of Iowa (Protocol 9052226) or the University of Minnesota (Protocol 2203-39886A). Mice were housed under standard conditions with access to standard chow. At the University of Iowa, the Cre/Lox system was used to generate CD4Cre+ RGS2flox/flox mice. C57BL/6J transgenic mice expressing Cre recombinase under the CD4 gene enhancer, promoter, and silencer sequence (Jackson Laboratories, Stock No. 017336) were crossed with B6SJLF1/J mice to create RGS2loxP/loxP mice, where exons 2–4 of the RGS2 gene were flanked by loxP sites (University of Iowa Genome Editing Core). CD4Cre+ mice were then crossed with RGS2flox/flox mice to generate CD4Cre+ RGS2flox/flox knockout (CD4 RGS2^KO^) mice. Littermate CD4Cre-RGS2flox/flox mice served as controls (CD4 RGS2^CTL^). Since these mice were backcrossed for only three generations to C57BL/6J, they were of mixed B6SJLF1/J background and are referred to as non-congenic CD4 RGS2 mice. At the University of Minnesota, CD4Cre (Stock No. 017336) and RGS2loxP/loxP (Stock No. 037619) mice, fully backcrossed onto the C57BL/6J background, were obtained from Jackson Laboratories. These congenic mice were crossed to generate CD4Cre+ RGS2flox/flox knockout (congenic CD4 RGS2^KO^) and littermate control (congenic CD4 RGS2^CTL^) mice. At the start of this study, non-congenic CD4 RGS2^KO^ and CD4 RGS2^CTL^ (University of Iowa) mice were used due to the extended time required to generate congenic strains, which typically involves backcrossing for at least 10 generations. The non-congenic animals were backcrossed for three generations to C57BL/6J, resulting in a mixed genetic background. Although congenic strains were eventually established and used for subsequent experiments, early studies were conducted with non-congenic mice to expedite initial data collection while the congenic lines were being developed. Genotyping for all mice was performed by Transnetyx (Cordova, TN).

### Superovulation Procedure

Four-week-old non-congenic or congenic female CD4 RGS2^KO^ and littermate control (CD4 RGS2^CTL^) mice were superovulated via intraperitoneal (IP) injection of 0.1 mL of 50 IU/mL pregnant mare serum gonadotropin (5.0 IU) at 1 pm on day 0. Ovulation was induced with an IP injection of 0.1 mL of 50 IU/mL human choriogonadotropin (hCG; 5.0 IU) 24 hours later on day 2. Ovaries and uteri were harvested approximately 19 hours after the hCG injection on day 3. Bilateral oviducts were dissected to isolate and count oocytes. Single-cell suspensions from the ovaries and uterus were prepared for flow cytometric analysis.

### Cell Preparation and Flow Cytometric Analysis

Single-cell suspensions were generated from the uterus and one ovary of each mouse via tissue digestion. Tissues were digested in media containing 1.5 mg/mL collagenase, 5% fetal bovine serum, Hank’s balanced salt solution, 10 nM HEPES, and DNase I, incubated at 37°C for 30 minutes with occasional vortexing. After digestion, tissues were mechanically dissociated and passed through a 70 µm strainer, followed by multiple washes. Cells were stained with fluorochrome-conjugated antibodies (Supplementary Table 1), and isotype control antibodies were used for background fluorescence. For intracellular staining of cytokines and transcription factors, the Intracellular Fixation and Permeabilization Buffer Set (Invitrogen, Waltham, MA) was used. Flow cytometric data were collected within 24 hours on a flow cytometer and analyzed with FlowJo software (Treestar Inc., Ashland, OR). Dead cells were excluded based on forward/side light scatter, and single cells were identified using forward scatter-area (FSC-A) vs. forward scatter-height (FSC-H). Gating was performed on CD45+ CD3+ CD4+ cells, with further analysis of CD4+ subsets, including T_H_1 (CD4+CCR5+; CD4+CXCR3+), T_H_2 (CD4+I L-4+; CD4+ IL-10+; CD4+ CXCR4+), T_H_17 (CD4+ CCR6+; CD4+ IL-17+), and Treg (CD4+ FOXP3+; CD4+CD25+; CD4+ LAP+). Macrophages were identified within the CD45+ CD3-population using markers F4/80+ CD11b+; F4/80+ CD86+; F4/80+ IL-12+; F4/80+ IL-1β+; F4/80+ iNOS+; and F4/80+ Arg1+. Natural killer (NK) cells were identified as CD45+ CD3-NKp46+ CD122+ and NKp46+ CD69+.

### Estradiol and Progesterone Quantification

Blood was collected at euthanasia from CD4 RGS2^KO^ and littermate control mice. After allowing the blood to clot for 30 minutes, samples were centrifuged at 4°C for 20 minutes at 1500 x g. Serum was stored at -80°C until ELISA assays were performed. Serum estradiol (Calbiotech, El Cajon, CA) and progesterone (Invitrogen, Waltham, MA) concentrations were measured using commercially available ELISA kits.

### Statistical Analysis

Statistical analyses for continuous variables were performed using two-tailed Student’s t-tests with unequal variance or two-way ANOVA with multiple comparisons, as appropriate (GraphPad Prism 8, La Jolla, CA). A p-value of α = 0.05 was considered statistically significant, or as determined by Bonferroni correction for multiple comparisons in ANOVA.

## Results

### Impact of Loss of RGS2 in CD4+ T Cells on Oocyte Numbers in Non-Congenic Mice

At the start of this study, non-congenic CD4 RGS2^KO^ and CD4 RGS2^CTL^ mice were used due to the time required to generate congenic strains, which involves at least 10 generations of backcrossing. We first aimed to investigate the effect that KO of RGS2 in CD4+ T cells had on follicular development and ovulation. To achieve this, we isolated and counted oocytes in prepubescent 4-week-old CD4 RGS2^KO^ and CD4 RGS2^CTL^ mice following superovulation. In non-congenic mice, the number of oocytes ovulated per cycle (KO 50 ± 9 vs. CTL 36 ± 10, p= 0.030) and per ovary (KO: 25 ± 5 vs. CTL: 17 ± 6, p= 0.030) were significantly higher in CD4 RGS2^KO^ mice compared to controls (Supplementary Table 2).

### Leukocyte Numbers and Frequencies in Non-Congenic CD4 RGS2^KO^ Mice

Loss of RGS2 in CD4+ T cells may affect both overall leukocyte numbers and the proportions of different leukocyte populations within the ovary and uterus. To investigate this possibility, we undertook flow cytometric analysis of total leukocytes as well as leukocyte subsets. Flow cytometric analysis revealed that uterine total leukocyte counts (KO: 3.9 × 10□ ± 6.9 × 10□ vs. CTL: 2.6 × 10□ ± 9.2 × 10□, p=0.0149; Supplementary Table 3) and non-T cells (CD3-) (KO: 3500 ± 1109.9 vs. CTL: 2109.1 ± 222.6, p= 0.0008; Supplementary Table 6) were significantly elevated in CD4 RGS2^KO^ mice. However, RGS2 knockout in CD4+ T cells did not affect CD4+ T_H_ subset total numbers or proportions in the ovary (Supplementary Tables 4 and 5) or uterus (Supplementary Tables 6 and 7). Ovarian and uterine macrophage and NK cell numbers and frequencies were also not altered in non-congenic CD4 RGS2^KO^ mice (Supplementary Tables 4-7).

### Congenic CD4 RGS2^KO^ Oocyte Numbers

Baseline immune cell populations and oocyte numbers can vary significantly between different mouse strains, complicating the comparison of these parameters in our initial experiments. The use of mice of a mixed genetic background (non-congenic) may influence the analysis of oocyte numbers and leukocyte populations, potentially introducing variability that could affect the interpretation of results. To account for these variables, we performed additional experiments on congenic (fully backcrossed) mice. As observed in non-congenic mice (Supplementary Table 2), loss of RGS2 specifically in CD4+ T cells resulted in significantly more oocytes (Figure 1).

**Figure 1:**
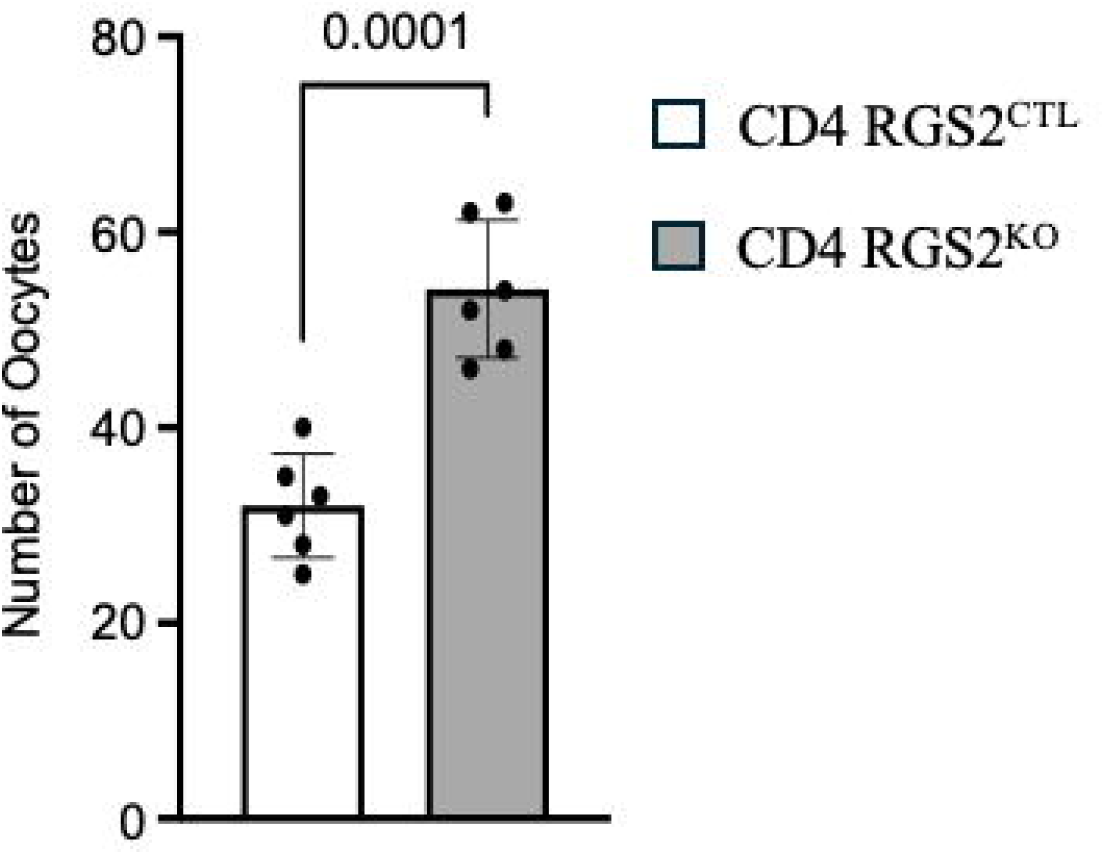
Increased Oocyte Numbers in Congenic CD4 RGS2 KO Mice. Data are mean ± SEM.

### Congenic CD4 RGS2^KO^ Leukocyte Numbers and T_H_ Cell Frequencies

While the number of total leukocytes in the ovaries were comparable between CD4 RGS2^CTL^ and CD4 RGS2^KO^ mice (Figure 2A), the number of leukocytes in the uterus of CD4 RGS2^KO^ mice was significantly increased compared to CD4 RGS2^CTL^ (Figure 2B). We next evaluated the frequencies of T_H_ cells within the ovary and uterus. Pro-inflammatory T_H_ 1 and T_H_17 cell frequencies were similar between the CD4 RGS2^KO^ and CD4 RGS2^CTL^ in the ovary (Figure 3A) and uterus (Figure 3C). Both anti-inflammatory Treg and T_H_2 were elevated in the ovary of CD4 RGS2^KO^ mice (Figure 3B), while only T_H_2 was elevated in the uterus of CD4 RGS2^KO^ mice (Figure 3D) compared to CD4 RGS2^CTL^ mice.

**Figure 2:**
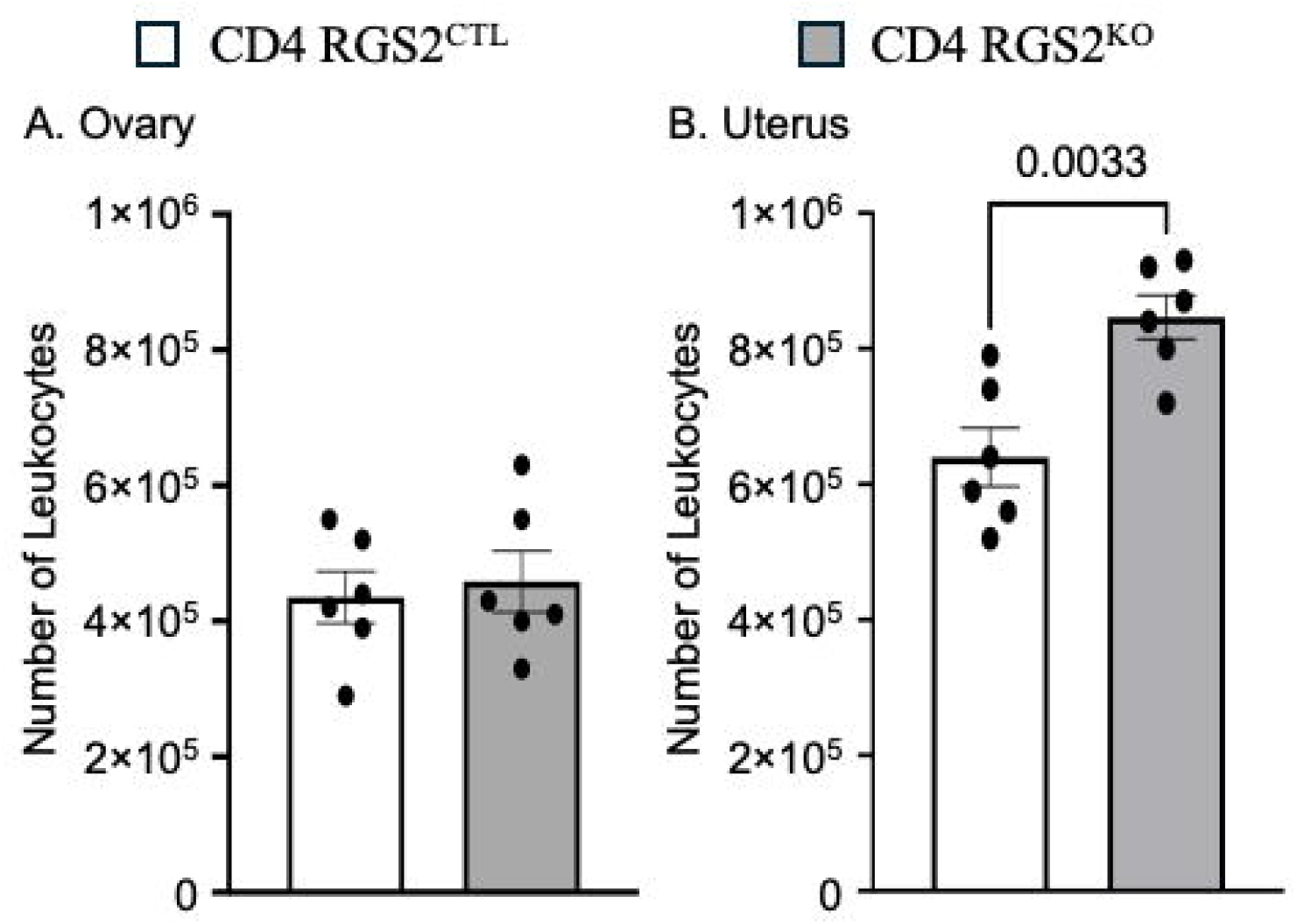
Increased Uterine Total Leukocyte Numbers in Congenic CD4 RGS2KO Mice. Number of leukocytes in the (A) ovary and (B) uterus in CD4 RGS2 KO and control mice. Data are mean ± SEM. ^*^=p<0.05.

**Figure 3.**
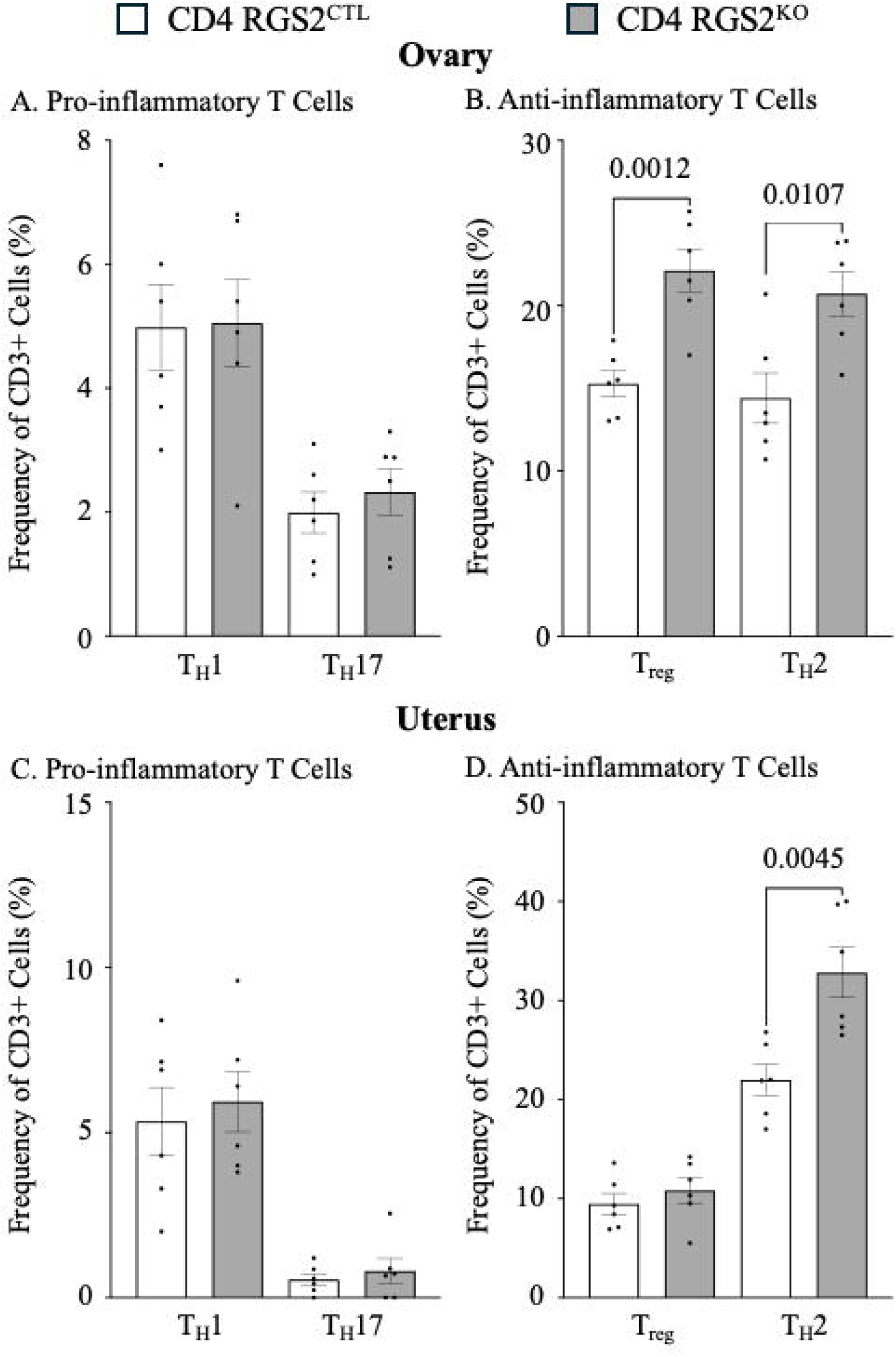
Increased Frequency of Ovarian and Uterine Anti-Inflammatory T Cells in CD4 RGS2 KO Congenic Mice. (A) Frequency of pro-inflammatory T_H_1 and T_H_17 in the ovary of congenic mice. (B) Frequency of anti-inflammatory T_reg_ and T_H_2 cells in the ovary of congenic mice. (C) Frequency of pro-inflammatory T_H_1 and T_H_17 in the uterus of congenic mice. (D) Frequency of anti-inflammatory T_reg_ and T_H_2 cells in the uterus of congenic mice. Data are mean ± SEM. *=p<0.05.

### Estradiol and Progesterone

We evaluated serum estradiol and progesterone in the mice at the time of euthanasia. Both estradiol (Figure 4A) and progesterone (Figure 4B) serum concentrations were elevated in CD4 RGS2^KO^ mice compared to CD4 RGS2^CTL^ mice.

**Figure 4.**
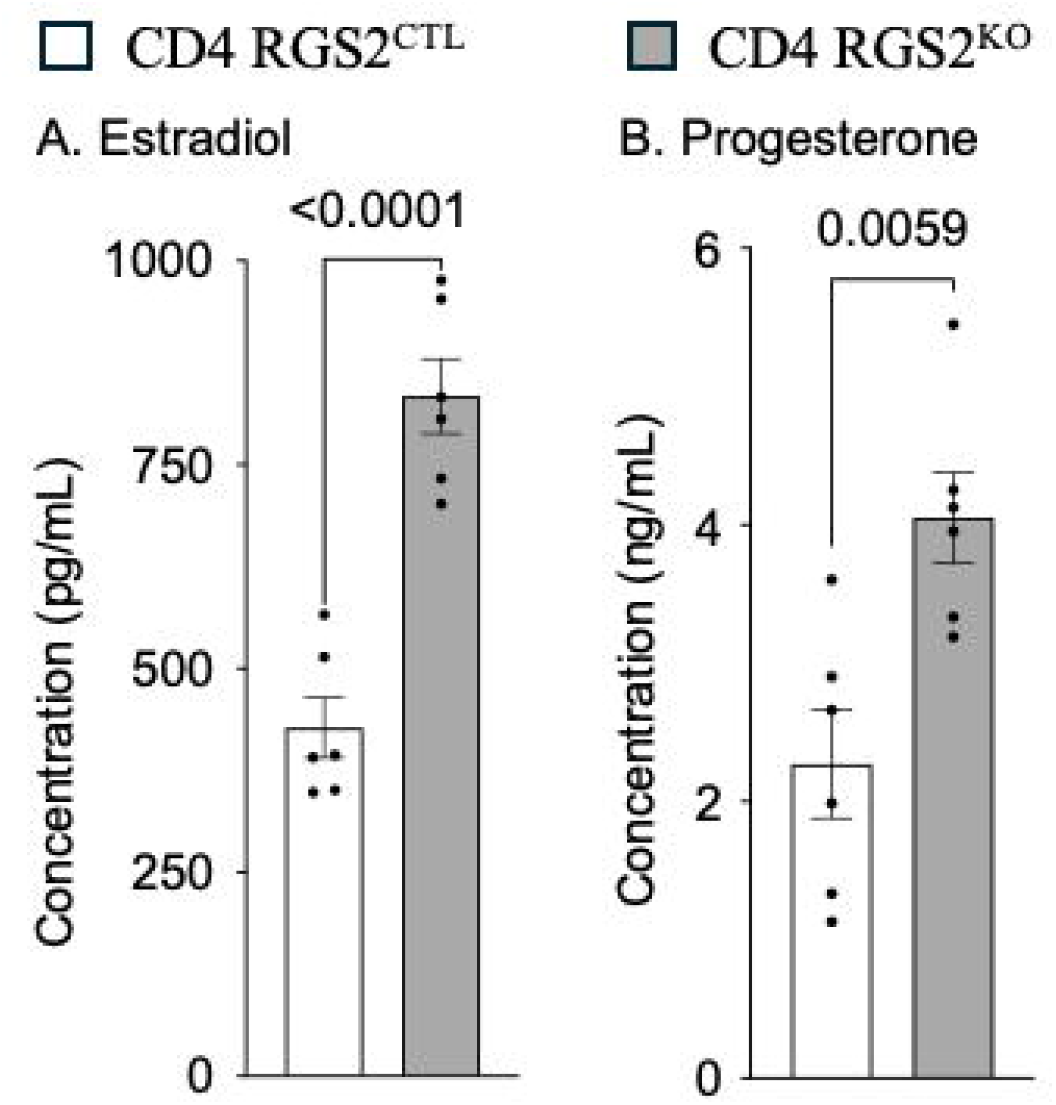
Increased Circulating Estradiol and Progesterone in Congenic CD4 RGS2 KO mice. Concentration of (A) estradiol and (B) progesterone in the serum of congenic mice. Data are mean ± SEM. *=p<0.05.

## Discussion

In this study, we investigated the consequences of knockout of RGS2 in CD4+ T cells on ovarian and uterine function. Our hypothesis was that the absence of RGS2 would reduce inflammatory T_H_ cell populations in reproductive tissues, impair ovulation, and decrease oocyte yield. At the start of this study, non-congenic CD4 RGS2^KO^ and CD4 RGS2^CTL^ mice were used due to the time required to generate congenic strains, which involves at least 10 generations of backcrossing. The results showed that non-congenic CD4 RGS2^KO^ mice exhibited a significant increase in the number of oocytes ovulated per cycle and per ovary compared to controls, suggesting that the presence of RGS2 in CD4+ T cells may negatively regulate oocyte yield, potentially through its effects on immune cell populations.

We also assessed leukocyte populations within the ovaries and uteri of non-congenic CD4 RGS2^KO^ mice. Our findings in non-congenic mice indicated that the total leukocyte counts, as well as the proportion of CD3-cells (non-T cell leukocytes), were significantly elevated in the KO group. However, RGS2 loss did not affect the total numbers or proportions of CD4+ T_H_ cell subsets, nor did it alter the populations of macrophages or NK cells in the ovaries and uteri. These initial findings suggested that while RGS2 in CD4+ T cells may modulate overall immune populations, its absence did not significantly disrupt the differentiation or activation of specific immune cell subsets. Baseline immune cell populations and oocyte numbers can vary dramatically between different mouse strains, which may further complicate the analysis when using mice of a mixed genetic background. Given that these mice were non-congenic, with a mixed genetic background that could introduce variability, we subsequently employed congenic mice to further evaluate the impact of RGS2 loss in CD4+ T cells on leukocyte populations and reproductive function.

To further investigate these effects, we performed experiments with congenic CD4 RGS2^KO^ and CD4 RGS2^CTL^ mice. Using congenic mice, which were fully backcrossed onto the C57BL/6J background, allowed us to reduce genetic variability and provide more reliable comparisons. The results mirrored those from the non-congenic mice, confirming that RGS2 loss in CD4+ T cells led to increased oocyte production. Unlike in the non-congenic CD4 RGS2 mice, we observed significant increases in anti-inflammatory T_H_2 and Treg cells within the CD4 RGS2^KO^ mice. This impact on oocyte production aligns with previous research indicating that RGS2 also plays a critical role in regulating oocyte maturation and function, as it suppresses premature calcium release in oocytes, preventing parthenogenetic activation prior to fertilization [1-4] [6-9]. Moreover, RGS2 expression has been shown to be essential for proper meiotic spindle formation in oocytes [7], and its role as a gene marker associated with IVF outcomes underscores its importance in reproductive biology [14].

Furthermore, the analysis of estradiol and progesterone levels revealed that both hormones were significantly elevated in CD4 RGS2^KO^ mice compared to controls. Elevated estradiol and progesterone levels are consistent with the observed increase in oocyte yield, as these hormones are produced by granulosa cells and corpora lutea respectively. Reproductive hormones including estrogen and progesterone are known to influence T_H_ cell populations [15-17]. High levels of estradiol have been shown to increase T_H_2 and Treg cells, while decreasing T_H_1 and T_H_17 cells [15], thus creating a more anti-inflammatory environment. Our findings indicate that the loss of RGS2 in CD4+ T cells may also have implications on the hormonal environment during the reproductive cycle which can impact downstream events important for reproduction.

Immune cells play vital roles in the processes leading to successful ovulation and endometrial development through the secretion of cytokines and chemokines that promote angiogenesis and tissue remodeling [1-4]. While we did not observe significant alterations in CD4+ T cell populations in the ovary, the elevated total uterine lymphocyte count and CD3-cell populations suggest that RGS2 loss may affect immune function in these tissues. Cytokines like TNF-α, IL-1, IL-6, and IL-8 have been shown to influence follicular growth and steroidogenesis, while also interacting with hormones like FSH and LH, which are critical for ovulation and endometrial function [18-23]. Therefore, future studies should investigate the production and function of cytokines and chemokines secreted by lymphocytes within the ovary and uterus, as these factors may influence follicular recruitment, oocyte numbers at the time of ovulation, and endometrial function.

## Conclusion

Our study highlights the important role of RGS2 in CD4+ T cells within the context of reproduction. The dysregulation of immune responses due to RGS2 knockout appears to enhance oocyte production while leaving the differentiation and activation of many immune subsets relatively unaffected. Further research is warranted to elucidate the precise mechanisms by which RGS2 influences reproductive outcomes, including its impact on endometrial receptivity and successful implantation. This work contributes to a growing understanding of the interplay between the immune system and reproductive health, which could ultimately aid in identifying novel therapeutic targets for unexplained infertility.

## Acknowledgments

The authors would like to thank Dr. Mark Santillan for providing the non-congenic breeding pair that initiated this study.

---

**Supplementary Figure 1:**
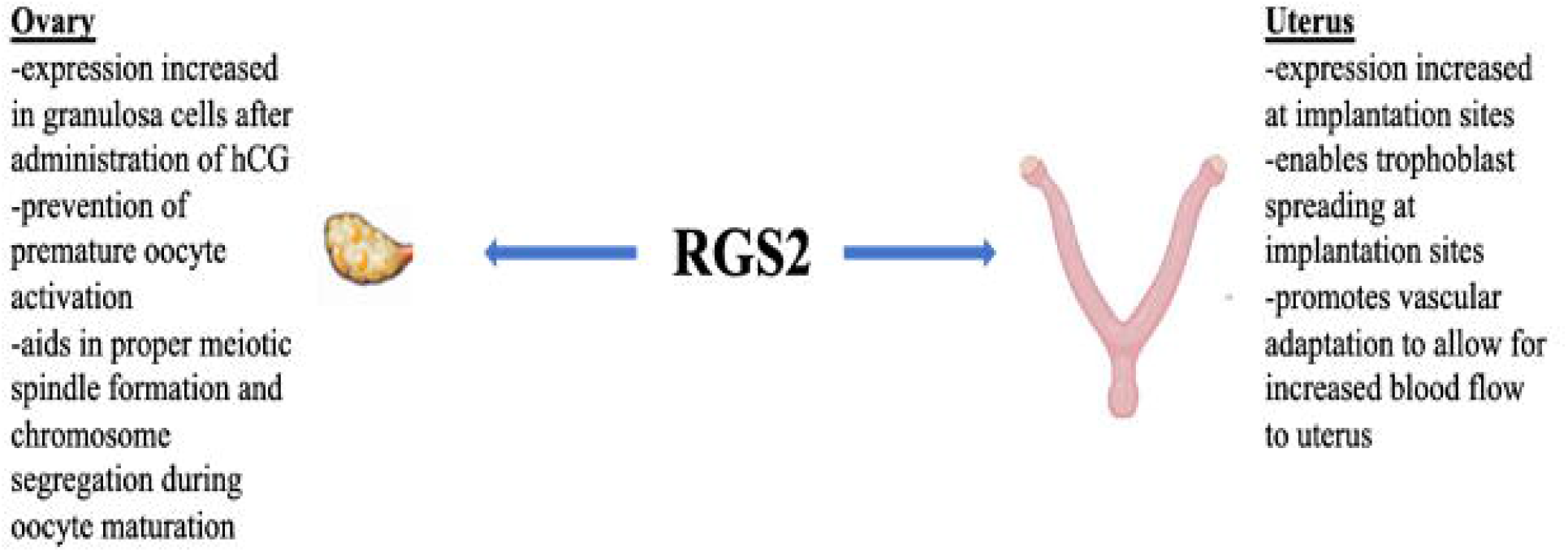
RGS2 Function in the Ovary and Uterus

## References

1. Duffy, D.M., et al., Ovulation: Parallels With Inflammatory Processes. Endocr Rev, 2019. 40: p. 369–416.

2. Bernhardt, M.L., et al., Regulator of G-protein signaling 2 (RGS2) suppresses premature calcium release in mouse eggs. Development, 2015. 142: p. 2633–40.

3. Ye, H., et al., The effect of the immune system on ovarian function and features of ovarian germline stem cells. Springerplus, 2016. 5: p. 990.

4. Jiang, M., et al., Uterine RGS2 expression is regulated by exogenous estrogen and progesterone in ovariectomized mice, and downregulation of RGS2 expression in artificial decidualized ESCs inhibits trophoblast spreading in vitro. Mol Reprod Dev, 2019. 86: p. 88–99.

5. Druey, K.M., Emerging Roles of Regulators of G Protein Signaling (RGS) Proteins in the Immune System. Adv Immunol, 2017. 136: p. 315–351.

6. Xie, Z., E.C. Chan, and K.M. Druey, R4 Regulator of G Protein Signaling (RGS) Proteins in Inflammation and Immunity. AAPS J, 2016. 18: p. 294–304.

7. Jiang, M.X., et al., Inhibition of the Binding between RGS2 and beta-Tubulin Interferes with Spindle Formation and Chromosome Segregation during Mouse Oocyte Maturation In Vitro. PLoS One, 2016. 11: p. e0159535.

8. Koch, J.N., et al., Regulator of G Protein Signaling 2 Facilitates Uterine Artery Adaptation During Pregnancy in Mice. J Am Heart Assoc, 2019. 8(9): p. e010917.

9. Jie, L., et al., RGS2 squelches vascular Gi/o and Gq signaling to modulate myogenic tone and promote uterine blood flow. Physiol Rep, 2016. 4.

10. Perschbacher, K.J., et al., Reduced mRNA Expression of RGS2 (Regulator of G Protein Signaling-2) in the Placenta Is Associated With Human Preeclampsia and Sufficient to Cause Features of the Disorder in Mice. Hypertension, 2020. 75: p. 569–579.

11. Feuerstein, P., et al., Genomic assessment of human cumulus cell marker genes as predictors of oocyte developmental competence: impact of various experimental factors. PLoS One, 2012. 7: p. e40449.

12. Suzuki, T., et al., Leukocytes in normal-cycling human ovaries: immunohistochemical distribution and characterization. Hum Reprod, 1998. 13: p. 2186–91.

13. Jasper, M.J., K.P. Tremellen, and S.A. Robertson, Primary unexplained infertility is associated with reduced expression of the T-regulatory cell transcription factor Foxp3 in endometrial tissue. Mol Hum Reprod, 2006. 12: p. 301–8.

14. Hamel, M., et al., Genomic assessment of follicular marker genes as pregnancy predictors for human IVF. Mol Hum Reprod, 2010. 16: p. 87–96.

15. Alanazi, H., et al., The impact of reproductive hormones on T cell immunity; normal and assisted reproductive cycles. J Reprod Immunol, 2024. 165: p. 104295.

16. Papapavlou, G., et al., Differential effects of estradiol and progesterone on human T cell activation in vitro. Eur J Immunol, 2021. 51: p. 2430–2440.

17. Hu, C., et al., Supraphysiological estradiol promotes human T follicular helper cell differentiation and favours humoural immunity during in vitro fertilization. J Cell Mol Med, 2021. 25: p. 6524–6534.

18. Buyalos, R.P., J.M. Watson, and O. Martinez-Maza, Detection of interleukin-6 in human follicular fluid. Fertil Steril, 1992. 57: p. 1230–4.

19. Wang, L.J., et al., Localisation of mRNA for interleukin-1 receptor and interleukin-1 receptor antagonist in the rat ovary. J Endocrinol, 1997. 152: p. 11–7.

20. Wang, L.J., et al., Tumor necrosis factor alpha in the human ovary: presence in follicular fluid and effects on cell proliferation and prostaglandin production. Fertil Steril, 1992. 58: p. 934–40.

21. Imai, F., et al., IL-6 up-regulates the expression of rat LH receptors during granulosa cell differentiation. Endocrinology, 2014. 155: p. 1436–44.

22. Nakao, K., et al., TNF-alpha Suppressed FSH-Induced LH Receptor Expression Through Transcriptional Regulation in Rat Granulosa Cells. Endocrinology, 2015. 156: p. 3192–202.

23. Buscher, U., F.C.K Chen, H. Schmiady, <Cytokines in the follicular fluid of stimulated and non-stimulated human ovaries; is ovulation a suppressed inflammatory reaction.pdf>. Hum Reprod, 1999. 14: p. 162–166.

