## Supplemental Tables for "Regulator of G-Protein Signaling 2 Knockout in CD4+ T Cells Promotes Anti-Inflammatory T Cells, Enhancing Ovulation, and Oocyte Yield"

Supplementary Table 1: Flow Cytometric Antibodies

| Target | Fluorophore | Company | ID Number |
| --- | --- | --- | --- |
| Arg1 | PE | Invitrogen | 12-3697-82 |
| CCR5 | PE | Invitrogen | 12-1951-82 |
| CCR6 | eFluor660 | Invitrogen | 50-7196-82 |
| CD3 | eFluor450 | Invitrogen | 48-0031-82 |
| CD4 | PerCPeFluor710 | Invitrogen | 46-0042-82 |
| CD8 | A700 | Invitrogen | 56-0081-82 |
| CD11b | APC | Invitrogen | 17-0112-82 |
| CD25 | A488 | Invitrogen | 53-0252-82 |
| CD45 | SB600 | Invitrogen | 63-0451-82 |
| CD62L | SB702 | Invitrogen | 67-0621-82 |
| CD69 | Fitc | Invitrogen | 11-0691-82 |
| CD86 | APCeFluor780 | Invitrogen | 47-0862-82 |
| CD122 | PECy7 | Invitrogen | 25-1222-82 |
| CXCR3 | PECy7 | Invitrogen | 25-1831-82 |
| CXCR4 | APCeFluor780 | Invitrogen | 47-9991-80 |
| F4/80 | SB645 | Invitrogen | 64-4801-82 |
| FOXP3 | A700 | Invitrogen | 56-5773-82 |
| IL-1 β | PerCPeFluor710 | Invitrogen | 46-7114-82 |
| IL-4 | A488 | Invitrogen | 53-7041-82 |
| IL-10 | PE | Invitrogen | 12-7101-82 |
| IL-12 | PECy7 | Invitrogen | 25-7123-82 |
| IL-17 | A488 | Invitrogen | 53-7177-81 |
| LAP | PECy7 | Invitrogen | 25-9821-82 |
| NKp46 | PE | Invitrogen | 12-3351-82 |

Supplementary Table 2: Loss of RGS2 in CD4+ T Cells Increases Oocyte Numbers in Non-Congenic Mice

|  | CD4 RGS2 Control | CD4 RGS2 Knockout | p-value |
| --- | --- | --- | --- |
| Number of Oocytes | 36 ±10 (n=11) | 50 ±9 (n=4) | 0.030 |
| Number of Oocytes/Ovary | 17±6 (n=12) | 25 ±5 (n=4) | 0.030 |

Supplementary Table 3: Uterine Total Leukocyte Numbers are Increased in Non-Congenic CD4 RGS2 KO Mice

|  | CD4 RGS Control | CD4 RGS Knockout | p-value |
| --- | --- | --- | --- |
| Total Ovary Leukocytes | 8.9 x 10^4^ ± 3.1 x 10^4^ (n=12) | 1.0 x 10^4^ ±3 .9 x 10^4^ (n=5) | 0.4279 |
| Total Uterine Leukocytes | 2.6 x 10^4^ ± 9.2 x 10^4^ (n=12) | 3.9 x 10^4^ ± 6.9 x10^4^ (n=5) | 0.0149 |

Supplementary Table 4: Loss of RGS2 in CD4+ T Cells Does Not Alter Ovarian Leukocyte Populations in Non-Congenic Mice

|  | CD4 RGS2 Control | | | CD4 RGS2 Knockout | | |  |
| --- | --- | --- | --- | --- | --- | --- | --- |
| Cell Type | Mean | SEM | N | Mean | SEM | N | p-value |
| CD3- | 11150 | 2385.48 | 12 | 16860.00 | 6268.94 | 5 | 0.264 |
| CD3+ | 1076.67 | 202.05 | 12 | 900.00 | 307.75 | 5 | >0.999 |
| CD4+ | 482.25 | 122.68 | 12 | 526.60 | 287.57 | 5 | >0.999 |
| Treg | 42.33 | 11.58 | 9 | 63.40 | 37.52 | 5 | >0.999 |
| T_H_1 | 296.67 | 91.09 | 6 | 606.67 | 496.7 | 3 | >0.999 |
| T_H_17 | 4.00 | 2.58 | 7 | 0.00 | 0.00 | 3 | >0.999 |
| T_H_2 | 118.50 | 51.21 | 6 | 161.60 | 83.60 | 5 | >0.999 |
| NK Cell | 61.33 | 20.85 | 6 | 144.80 | 48.34 | 5 | >0.999 |
| NK Activated | 113.67 | 50.98 | 9 | 46.67 | 27.44 | 3 | >0.999 |
| Macrophage | 7600.00 | 1871.76 | 12 | 9248.00 | 5440.92 | 5 | >0.999 |
| M1 Macrophage | 243.00 | 212.57 | 6 | 258.00 | 202.00 | 2 | n/a* |
| M2 Macrophage | 823.33 | 603.78 | 3 | 12.00 | 12.00 | 2 | n/a* |

* n/a= due to limited sample size, statistical measures could not be performed

Supplementary Table 5: Maintained Ovarian Leukocyte Frequencies in CD4 RGS2 KO Non-Congenic Mice

|  | CD4 RGS2 Control | | | CD4 RGS2 Knockout | | |  |
| --- | --- | --- | --- | --- | --- | --- | --- |
| Cell Type | Mean | SEM | N | Mean | SEM | N | p-value |
| CD3- | 88.47 | 2.71 | 12 | 93.33 | 1.18 | 5 | >0.999 |
| CD3+ | 11.52 | 2.71 | 12 | 6.68 | 1.18 | 5 | >0.999 |
| CD4+ | 43.89 | 5.10 | 10 | 42.34 | 15.56 | 5 | >0.999 |
| Treg | 12.83 | 5.32 | 6 | 13.24 | 10.69 | 3 | >0.999 |
| T_H_1 | 60.98 | 11.96 | 6 | 49.33 | 25.36 | 3 | 0.9948 |
| T_H_17 | 0.695 | 0.69 | 6 | 0.00 | 0.00 | 3 | >0.999 |
| T_H_2 | 30.06 | 13.05 | 6 | 28.8 | 18.21 | 3 | >0.999 |
| NK Cell | 1.33 | 0.30 | 12 | 1.02 | 0.12 | 5 | >0.999 |
| NK Activated | 1.04 | 0.35 | 9 | 0.16 | 0.09 | 3 | >0.999 |
| Macrophage | 59.63 | 6.49 | 12 | 47.26 | 15.06 | 5 | 0.924 |
| M1 Macrophage | 4.97 | 3.89 | 6 | 12.52 | 10.08 | 2 | n/a* |
| M2 Macrophage | 16.92 | 12.92 | 3 | 0.59 | 0.59 | 2 | n/a* |

*n/a= due to limited sample size, statistical measures could not be performed

Supplementary Table 6: Loss of RGS2 in CD4+ T Cells Increases the Number of Uterine Non-T Cells in Non-Congenic Mice

|  | CD4 RGS2 Control | | | CD4 RGS2 Knockout | | |  |
| --- | --- | --- | --- | --- | --- | --- | --- |
| Cell Type | Mean | SEM | N | Mean | SEM | N | p-value |
| CD3- | 2109.09 | 222.57 | 11 | 3500.00 | 1109.96 | 5 | 0.0008 |
| CD3+ | 173.64 | 27.55 | 11 | 355.40 | 100.51 | 5 | >0.999 |
| CD4+ | 98.73 | 14.23 | 11 | 178.80 | 39.77 | 5 | >0.999 |
| Treg | 0.00 | 0.00 | 5 | 8.67 | 8.67 | 3 | >0.999 |
| T_H_1 | 87.80 | 30.03 | 5 | 151.67 | 51.67 | 3 | >0.999 |
| T_H_17 | 0.00 | 0.00 | 5 | 196.67 | 35.28 | 3 | >0.999 |
| T_H_2 | 0.00 | 0.00 | 5 | 36.67 | 36.67 | 3 | >0.999 |
| NK Cell | 47.00 | 8.34 | 6 | 69.33 | 35.88 | 3 | >0.999 |
| NK Activated | 14.13 | 7.29 | 8 | 19.67 | 19.67 | 3 | >0.999 |
| Macrophage | 921.82 | 210.06 | 11 | 1308.00 | 497.33 | 5 | 0.969 |
| M1 Macrophage | 37.00 | 20.22 | 6 | 85.00 | 85.00 | 2 | n/a* |
| M2 Macrophage | 220.00 | 220.00 | 3 | 55.00 | 17.00 | 2 | n/a* |

*n/a= due to limited sample size, statistical measures could not be performed

Supplementary Table 7: KO of RGS2 in CD4 T Cells Does Not Alter Uterine Leukocyte Proportions in Non-Congenic Mice

|  | CD4 RGS2 Control | | | CD4 RGS2 Knockout | | |  |
| --- | --- | --- | --- | --- | --- | --- | --- |
| Cell Type | Mean | SEM | N | Mean | SEM | N | p-value |
| CD3- | 92.52 | 1.14 | 11 | 90.02 | 2.60 | 5 | >0.999 |
| CD3+ | 8.16 | 1.32 | 11 | 9.98 | 2.59 | 5 | >0.999 |
| CD4+ | 59.18 | 5.16 | 11 | 59.98 | 11.65 | 5 | >0.999 |
| Treg | 0.00 | 0.00 | 5 | 3.03 | 3.03 | 3 | >0.999 |
| T_H_1 | 100.00 | 0.00 | 5 | 91.40 | 8.60 | 3 | 0.991 |
| T_H_17 | 0.00 | 0.00 | 5 | 1.52 | 1.52 | 3 | >0.999 |
| T_H_2 | 0.00 | 0.00 | 5 | 12.62 | 12.62 | 3 | 0.859 |
| NK Cell | 1.83 | 0.42 | 11 | 1.87 | 0.76 | 5 | >0.999 |
| NK Activated | 0.58 | 0.33 | 8 | 0.58 | 0.58 | 3 | >0.999 |
| Macrophage | 42.69 | 7.13 | 11 | 38.84 | 9.02 | 5 | >0.999 |
| M1 Macrophage | 2.23 | 1.58 | 6 | 3.66 | 3.66 | 2 | n/a* |
| M2 Macrophage | 12.17 | 12.17 | 3 | 3.91 | 2.34 | 2 | n/a* |

*n/a= due to limited sample size, statistical measures could not be performed
